# Aberrant Hippocampal Neurogenesis is A Conserved Response to Stroke in Mice: A Multi-Center Multimodel Study

**DOI:** 10.64898/2026.02.19.706917

**Authors:** Francisco J. De Castro-Millán, Sandra Vázquez-Reyes, Carolina Peña-Martínez, Adrián Rodríguez-Llave, Carlos Parra-Pérez, Carmen Nieto-Vaquero, Gaia Brezzo, Kristy A. Zera, Dana Straus, Jennifer E. Goertz, Sanna H. Loppi, Rachel R. Crumpacker, Jennifer B. Frye, Danielle A. Becktel, Claudia Dames, Daniel Berchtold, Jill H. Fowler, Andreas Meisel, Josef Anrather, Kristian P. Doyle, Stuart M. Allan, Marion S. Buckwalter, Barry W. McColl, Alicia García-Culebras, María Isabel Cuartero, María Ángeles Moro

**Author notes:** Correspondence to: María A. Moro, María I. Cuartero or Alicia García-Culebras. Equal contribution (Francisco J. De Castro-Millán, Sandra Vázquez-Reyes).

## Abstract

**Background:** Adult hippocampal neurogenesis is markedly altered after cerebral ischemia. Although stroke increases the production of newborn neurons, many of these cells display aberrant morphological and positional features that impair their functional integration and contribute to long-term cognitive decline. Given the clinical heterogeneity of ischemic stroke and the persistent translational failures of preclinical approaches relying on single-model studies it remains unknown whether post-stroke neurogenic alterations are conserved across different experimental paradigms. This study aimed to define common and model-specific features of hippocampal neurogenesis across complementary focal ischemia models.

**Methods:** We performed a multi-center, multimodel analysis within the STROKE-IMPaCT consortium using permanent and transient middle cerebral artery occlusion (MCAO) paradigms (MCAO via ligation or cauterization under normoxic (dMCAO) or hypoxic conditions (dMCAO+Hypoxia); and filament-based tMCAO across six international sites. Brains from adult C57BL/6J mice were collected 3 days, 7 days, or 2 months after ischemia, sham, or naive conditions. Hippocampal cell proliferation (Ki67) and neuroblast density (DCX) were quantified, and the morphological maturation of newborn neurons was evaluated using high-resolution analyses of dendritic architecture and somatodendritic polarity. All analyses were performed blind to experimental group.

**Results:** Across all stroke models, ischemia induced a robust bilateral increase in hippocampal cell proliferation, most pronounced at 3 days and still elevated at 7 days, with levels returning to baseline by 2 months. Neuroblast density was similarly increased at 7 days, particularly in the ipsilateral hippocampus, but normalized by 2 months. Despite recovery in cell number, long-term morphological analysis revealed a consistent reduction in apical dendrite length and a higher proportion of neurons exhibiting aberrant features including ectopic localization, multipolar or inverted polarity, and abnormal lateral growth across all models. These abnormalities were observed both when pooling data across sites and when analyzing each model or center individually.

**Conclusions:** Ischemia induces an early, transient increase in hippocampal neurogenesis across diverse stroke paradigms, but the newborn neurons generated after stroke consistently display maladaptive morphological features. These cross-model, cross-site abnormalities indicate that aberrant hippocampal neurogenesis represents a robust hallmark of post-stroke pathology within the investigated species, independent of ischemia type or surgical approach, despite known differences in the spatial distribution of primary injury across experimental stroke models. Our findings support the concept that maladaptive neurogenesis may contribute to chronic post-stroke cognitive impairment and underscore the need to consider the quality not only the quantity of newborn neurons when developing therapeutic strategies.

## INTRODUCTION

The adult hippocampus retains a remarkable capacity for structural plasticity through the continuous generation of new granule neurons in the dentate gyrus (DG), a process extensively characterized in experimental models and supported by accumulating evidence in the adult primate and human brain^1–3^. This process, termed adult hippocampal neurogenesis (AHN), originates from neural stem cells residing in the subgranular zone (SGZ) and culminates in the functional integration of newborn neurons into pre-existing hippocampal circuits. Adult-born granule neurons undergo a prolonged and tightly regulated maturation sequence that confers distinct physiological properties and contributes to circuit plasticity, pattern separation, and memory updating^3,4^. Perturbations of this process have been implicated in hippocampal dysfunction across a range of neurological and vascular conditions. Cerebral ischemia is a potent modulator of AHN. Experimental stroke robustly activates the hippocampal neurogenic niche, inducing a marked increase in progenitor proliferation and neuroblast production, a phenomenon first described more than two decades ago and consistently reproduced across ischemia models^5–8^. This AHN response, initially interpreted as an endogenous repair mechanism, does not necessarily translate into functional benefit. Instead, neurons generated in the post-ischemic hippocampus frequently display aberrant features, including ectopic positioning, inverted or multipolar somatodendritic orientation, and altered dendritic growth and connectivity^9–12^. These structural abnormalities suggest impaired neuronal integration and maladaptive circuit remodeling post-stroke.

Importantly, such maladaptive neurogenesis is not merely an epiphenomenon. Aberrant AHN has been causally linked to long-term hippocampus-dependent cognitive dysfunction in rodents, as suppression of post-stroke neurogenesis ameliorates memory deficits, whereas further enhancement of neurogenesis exacerbates them^10,11^.

Together, these studies identify AHN as a critical mechanistic substrate through which ischemic injury can reshape hippocampal circuitry and influence cognitive outcome in experimental settings, even when the primary lesion is anatomically remote from the hippocampus.

Stroke in humans encompasses a wide spectrum of underlying pathophysiology, including large-artery occlusion, small-vessel disease, cardioembolism, microinfarction, and chronic hypoperfusion, each associated with distinct vascular, inflammatory, and metabolic environments. This heterogeneity strongly suggests that ischemia-induced hippocampal remodeling may not follow a single biological trajectory. Nevertheless, most preclinical studies of post-stroke AHN rely on individual models, and are conducted within single laboratories. Because of this, these studies differ substantially in lesion characteristics, severity, timing, markers, and analytical pipelines, leaving unresolved whether aberrant AHN represents a conserved response to cerebral ischemia or a model-dependent phenomenon.

Addressing this question requires a systematic approach capable of capturing the diversity of ischemic pathophysiology encountered clinically while minimizing laboratory-specific bias. To this end, the present study leverages a multi-center, multi-model design within the Stroke-IMPaCT consortium, a coordinated transatlantic network of six laboratories across Europe and North America applying distinct ischemic paradigms in parallel to enhance reproducibility, enable direct comparison across models, and increase the generalizability of mechanistic conclusions. Previous work within this consortium has highlighted how heterogeneous lesion profiles influence long-term functional outcomes and their detectability, underscoring the importance of cross-model analyses^13^.

Building on this, we systematically examine the hippocampal neurogenic response across multiple widely used stroke models, focusing on SGZ proliferation, neuroblast distribution, and the morphological maturation of adult-born granule neurons. By directly comparing these parameters across complementary ischemia paradigms, we aim to determine whether maladaptive hippocampal neurogenesis constitutes a conserved hallmark of cerebral ischemia. Identifying neurogenic features that remain stable across heterogeneous experimental designs is a critical step toward defining robust mechanisms of ischemia-induced hippocampal remodeling with higher translational relevance.

## METHODS

### Animals

Adult male C57BL/6J mice (12–15 weeks old) were used throughout the study. Animals were housed under a 12-hour light/dark cycle with *ad libitum* access to food and water. All procedures adhered to the ARRIVE (Animal Research: Reporting of In Vivo Experiments) guidelines and were approved by the corresponding institutional animal care and use committees (Centro Nacional de Investigaciones Cardiovasculares (CNIC): Animal Welfare Committee and Ministry of Environment and Territorial Planning of the Community of Madrid, RD 53/2013; PROEX 047/16, in compliance with European directives 86/609/EEC and 2010/63/EU; Charité Universitätsmedizin Berlin (CUB): Landesamt für Gesundheit und Soziales, Berlin, G0312/16 and G0167/20; UoE: Bioresearch and Veterinary Services Committee; University of Arizona (UoA), Stanford University (SU), Weill Cornell Medicine (WC): Institutional Animal Care and Use Committees). Experiments were conducted in accordance with relevant national and institutional regulations (European Directive 2010/63/EU; UK Animals Scientific Procedures Act 1986; NIH and US Public Health Service Guidelines). Despite ARRIVE and STEPS recommendations encouraging the inclusion of both sexes, only male mice were used due to the large sample size required in a multi-center design and the time-intensive nature of neurogenesis and morphometric analyses, which made the inclusion of both sexes incompatible with achieving balanced and adequately powered groups across all sites. Sham-operated and naive mice served as control groups. Animals were randomly assigned to experimental conditions, and investigators were blinded during tissue processing, imaging, and quantitative analyses.

Due to the extensive workload associated with tissue processing, neurogenesis quantification, and morphological assessments, only a subset of animals from the original cohort was selected for the present analyses. Animals were randomly selected from each experimental group and center to ensure representative sampling and minimize selection bias, based on predefined technical inclusion criteria. Specifically, mice were randomly chosen within each experimental group provided that (i) the presence and extent of ischemic injury were consistent with the expected model and time point, and (ii) tissue quality was adequate, with well-preserved morphology and no evident damage resulting from tissue processing^13^. The final number of mice included in the neurogenesis dataset was 284, distributed across time points: 3 days post-surgery, n=91 mice (20 naive, 35 sham, and 36 ischemic); 7 days post-surgery, n=92 mice (20 naive, 36 sham, and 36 ischemic); and 2 months post-surgery, n=101 (35 naive, 34 sham, and 32 ischemic). Detailed information on site-specific differences in mouse origin, housing, enrichment, and diet is provided in the Supplemental Material.

### Stroke Models

All surgeries were performed under inhalational anesthesia, and core temperature was maintained at 37°C throughout the procedure. Pre- and postoperative analgesia, as well as topical local anesthetics, were administered according to institutional protocols. Focal cerebral ischemia was induced using three established middle cerebral artery occlusion (MCAO) paradigms across the six participating centers^10,11,14–20^ (**Figure 1**). Permanent distal MCA occlusion (dMCAO) was implemented by distal MCA ligation at CNIC (Madrid) and by distal MCA cauterization at UoE (Edinburgh). A variant combining permanent MCA occlusion with systemic hypoxia (dMCAO + Hypoxia) was performed at UoA (Arizona) and SU (Stanford). Proximal transient MCA occlusion (tMCAO) using the intraluminal filament technique was carried out at CUB (Berlin) and WC (New York). For each model, littermate controls underwent the corresponding sham procedure, and naive mice served as non-surgical controls. Detailed methodology for each stroke model is described in the Supplemental Material.

**Figure 1.**
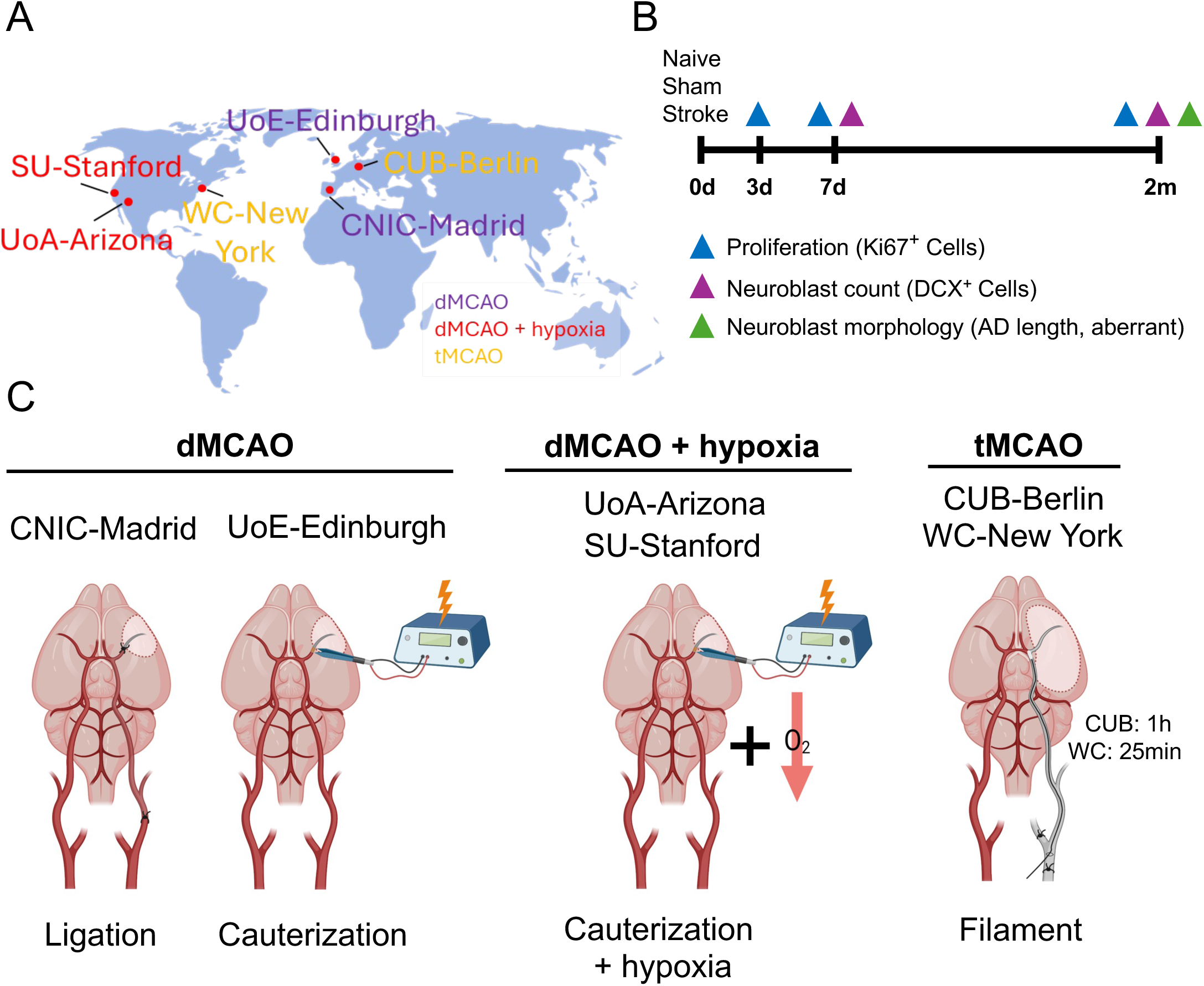
Multi-center experimental design and sampling strategy. (A) Geographic distribution of the six participating centers: CNIC–Madrid, UoE–Edinburgh, SU–Stanford, UoA–Arizona, WC–New York, and CUB–Berlin, together with the ischemia model used at each site. CNIC–Madrid and UoE–Edinburgh performed permanent distal MCA occlusion (dMCAO) via distal MCA ligation (Madrid) or cauterization (Edinburgh). SU–Stanford and UoA–Arizona applied a permanent distal ischemia paradigm based on distal MCA cauterization combined with systemic hypoxia (dMCAO + hypoxia). WC–New York and CUB–Berlin used transient filament-based proximal MCA occlusion (tMCAO). (B) Experimental timeline showing collection of naive, sham, and stroke samples at 3 days, 7 days, and 2 months post-stroke. Proliferation (Ki67⁺ cells) and neuroblast number (DCX⁺ cells) were quantified at 3 days and 7 days, while neuroblast morphology (apical dendrite length and aberrant phenotypes) was assessed at 2 months. All samples were processed using harmonized pipelines and analyzed under blinded conditions across centers. (C) Schematic representation of the surgical procedures used across centers. Within the dMCAO group, CNIC–Madrid performed distal MCA ligation, and UoE–Edinburgh applied distal MCA cauterization. Permanent distal ischemia combined with systemic hypoxia (dMCAO + hypoxia) was performed at SU–Stanford and UoA–Arizona using distal MCA cauterization followed by a controlled hypoxic period. Transient proximal occlusion (tMCAO) was carried out at CUB–Berlin (60 min) and WC–New York (25 min) using an intraluminal filament. Illustrations depict the vascular territory and technical approach specific to each paradigm. All schematic illustrations in panel C were created with BioRender.

### Tissue Collection and Processing

Mice were euthanized at 3 days, 7 days, or approximately 2 months after stroke by exsanguination and intracardiac perfusion with 0.9% saline (without anticoagulant) under deep anesthesia. Anesthetic regimens varied across centers and included isoflurane (UoE, UoA), sevoflurane (CNIC), sodium pentobarbital (WC), or xylazine/ketamine (SU, CUB). Following perfusion, brain tissue and blood samples were collected. Brains were removed, post-fixed in 4% paraformaldehyde in phosphate buffer for 24h and subsequently transferred to a 30% sucrose solution containing 0.1% sodium azide for cryoprotection. For long-term storage, brains were rapidly frozen by immersion in pre-chilled 2-methylbutane (Supelco) for approximately 10s at −80°C and kept at this temperature until sectioning.

All brain tissue was sectioned at 40μm thickness using a freezing microtome (UoA). Sections were sequentially collected into 16 tubes containing cryoprotective medium (30% glycerin, 30% ethylene glycol, 40% 0.5 M sodium phosphate buffer). All samples were stored at -20°C in cryoprotective solution until further processing.

### Histopathology and Immunostaining

#### Hematoxylin & Eosin (H&E) Staining

H&E staining was performed at the University of Edinburgh (UoE) following standard procedures using commercially available reagents (Instant Hematoxylin, Epredia; Eosin Y, Cell Path). Brain sections were washed in PBS at room temperature (RT) to remove cryoprotectant, mounted on gelatin-coated slides, rinsed in distilled water, and air-dried overnight prior to staining. Slides were processed as previously described^13^. Briefly, sections were dipped 10 times in 95% ethanol and rinsed in distilled water for 30s. Slides were incubated in hematoxylin for 90 s, rinsed in distilled water until clear, dipped in acid alcohol for 30 s, and rinsed again for 30s. Differentiation was performed in Scott’s tap water for 60 s. After a brief rinse in distilled water, slides were dipped once in eosin, washed, dehydrated through ascending ethanol solutions, cleared in xylene, and mounted with DPX. H&E staining was performed on two coronal sections per animal: one rostral (Bregma −0.70 to −0.82 mm) and one caudal (Bregma −1.70 to −2.18 mm).

In the 2 months post-surgery group, an analysis of the GCL (Granular cell layer) area was performed using digitized images of H&E staining samples. Animals with well-preserved and mounted sections containing hippocampal tissue were selected. The GCL area was measured using the “polygon selections” tool of ImageJ-Fiji (v1.54p) software and averaged across the different hippocampal sections from each animal.

### Immunofluorescence Staining

Immunofluorescence was performed on free-floating coronal hippocampal sections. Sections were washed three times in 0.1M PBS to remove cryoprotectant and incubated for 1h at RT in blocking/permeabilization solution containing 5% bovine serum albumin (BSA; Sigma-Aldrich) and 0.25% Triton X-100 (VWR Chemicals) in PBS, with gentle agitation.

Primary antibodies were diluted in the same blocking solution and applied overnight at 4°C with gentle shaking. The following primary antibodies were used: anti-DCX (rabbit, 1:250; Abcam), anti-Ki67 (rat, 1:200; Invitrogen). The next day, sections were washed three times (10 min each) in PBS containing 0.25% Triton X-100. Secondary antibodies were applied for 2h at RT. For DCX staining, a biotinylated anti-rabbit secondary antibody (goat, 1:250; BioNova) was followed by 1h incubation with streptavidin-488 (Invitrogen). Sections were counterstained with DAPI (1:5000, Merck) for 5 min, washed in PBS, and mounted on gelatin-coated slides using Aqua-Poly Mount (Polysciences). Mounted slides dried overnight at RT and were stored at 4°C until imaging.

### Confocal Imaging

Confocal images were acquired using a Leica SP8 microscope equipped with hybrid spectral detectors and a 40× oil-immersion objective (CNIC Microscopy Unit). Z-stack images were collected from at least three dentate gyrus sections per animal and processed in 3D using IMARIS software (v9.1.2, Bitplane).

### Quantifications of Different Cell Populations

The number of Ki67^+^ and DCX^+^ cells were quantified in the SGZ (Subgranular Zone) and GCL of the dentate gyrus. For some aberrant features like ectopic locations, those DCX^+^ cells located in the hilus or in the molecular layer were also considered. In sham-operated and naïve animals, ipsilateral and contralateral hemispheres could not be reliably distinguished; therefore, neurogenesis was quantified in both hemispheres, and the mean value was used for analysis. In ischemic animals, ipsilateral and contralateral hemispheres were quantified separately. The counting of the positive cells was carried out manually. Data of Ki67^+^ and DCX^+^ cells are expressed in number of cells per mm of DG (Dentate Gyrus), since the length of the DG was quantified with the “segmented lines” tool of ImageJ-Fiji software following the line between the SGZ and the hilus.

### Morphological Analysis of Neuronal Populations

Morphological analyses were performed in IMARIS using the Filaments tool. Measurements included apical dendrite (AD) length. Only cells whose soma and dendritic arbor were fully represented within the tissue section, and without overlapping with neighboring cells, were included in the analysis. Apical dendrite length was defined as the dendritic extension emerging from the soma and projecting radially through the granule cell layer until the first major bifurcation and was measured with the filaments tool of IMARIS software. Aberrant neuroblasts, including multipolar, inverted, lateral-growth, and ectopic (hilus or molecular layer) phenotypes, were quantified relative to the total number of DCX-positive cells. Due to low frequency, ectopic neurons in the hilus and molecular layer were pooled as a single ectopic category.

### Statistical Analysis

Statistical analyses were performed using Prism 10 (GraphPad Software Inc.). Prior to applying any statistical test, data were assessed for normality using Shapiro–Wilk test. If the data met the assumption of normality, parametric tests were applied. In cases where data did not follow a normal distribution, or when the sample size was n<6, non-parametric alternatives were employed. For comparisons between two groups, normally distributed data were analyzed with an unpaired t-test, while non-normally distributed data were compared using the Mann–Whitney U test. For comparisons among more than two groups with a single independent variable, one-way ANOVA was used if assumptions were met; otherwise, the Kruskal–Wallis test was applied, followed by Dunn’s post hoc test for pairwise comparisons. For analyses involving two independent variables, a two-way ANOVA was performed. For post hoc comparisons between groups, we applied Emmeans, Kruskal–Wallis tests with Dunn’s correction and, Mann–Whitney U tests with Holm–Šídák correction. All statistical analyses included corrections for multiple comparisons using statistical hypothesis testing. All statistical data were represented as the mean ± SEM, and in all studies, tests were considered significant when p<0.05. Excluded values from the study were identified and removed using various outlier detection criteria. The Grubbs’ test (α=0.05) was applied to exclude extreme values when only one value was considered an outlier. Additionally, the ROUT method (Q=1%) was used, and in certain cases, exclusion was based on the interquartile range (IQR), excluding values lower or higher than 1.5 times the IQR when the sample size was between 3 and 5.

## RESULTS

### STROKE INDUCES A BILATERAL INCREASE IN HIPPOCAMPAL PROLIFERATION ACROSS ALL STROKE MODELS

To characterize the hippocampal neurogenic response after stroke, we evaluated cell proliferation and neuroblast generation at defined post-ischemic time points. Proliferation was quantified by Ki67 immunostaining at 3 days, 7 days, and 2 months, whereas neuroblast numbers were assessed using doublecortin (DCX) at 7 days and 2 months **(Figure 1).** At the 2-month time point, we additionally examined the morphological features of newly generated neurons to determine whether long-term maturation was altered by ischemia. This multicenter study enabled the inclusion of several variants of permanent and transient middle cerebral artery occlusion (MCAO) in the analysis. The Madrid and Edinburgh groups employed permanent distal MCA occlusion (dMCAO) using distal MCA ligation (Madrid) or distal MCA cauterization (Edinburgh). The Stanford and Arizona groups applied a permanent distal ischemia paradigm based on distal MCA cauterization combined with a controlled hypoxic period (dMCAO + hypoxia). In contrast, the Berlin and New York groups used proximal intraluminal approaches, specifically transient MCAO (tMCAO) **(Figure 1).** We first quantified Ki67-positive cells in the SGZ of the DG across all experimental groups **(Figure 2)**. When data from all centers and stroke models were combined, a clear proliferative response emerged at both 3 and 7 days after ischemia **(Figure 2A-B)**. This increase was bilateral, affecting both the ipsilateral and contralateral hippocampus to a similar extent, and was more pronounced at 3 days, suggesting a rapid and early surge in cell-cycle re-entry following stroke. By 7 days, proliferation remained elevated but was generally reduced compared with the peak observed at 3 days. In contrast, no significant differences in the numbers of Ki67-positive cells were detected at the 2-month time point, indicating that the post-stroke proliferative response had returned to baseline. Notably, Ki67-positive cell numbers at 2 months were generally lower across all experimental groups, including naive animals, consistent with an age-dependent decline in baseline hippocampal proliferation. The early post-stroke increase in Ki67-positive cells was significant when compared with both sham-operated and naive mice, demonstrating that the proliferative response was specifically driven by the ischemic insult. These effects were consistently observed across the three major ischemic paradigms, dMCAO, dMCAO combined with hypoxia, and tMCAO **(Figure 2C)**, and were also evident when each site was analyzed individually **(Figure 2D-F)**.

**Figure 2.**
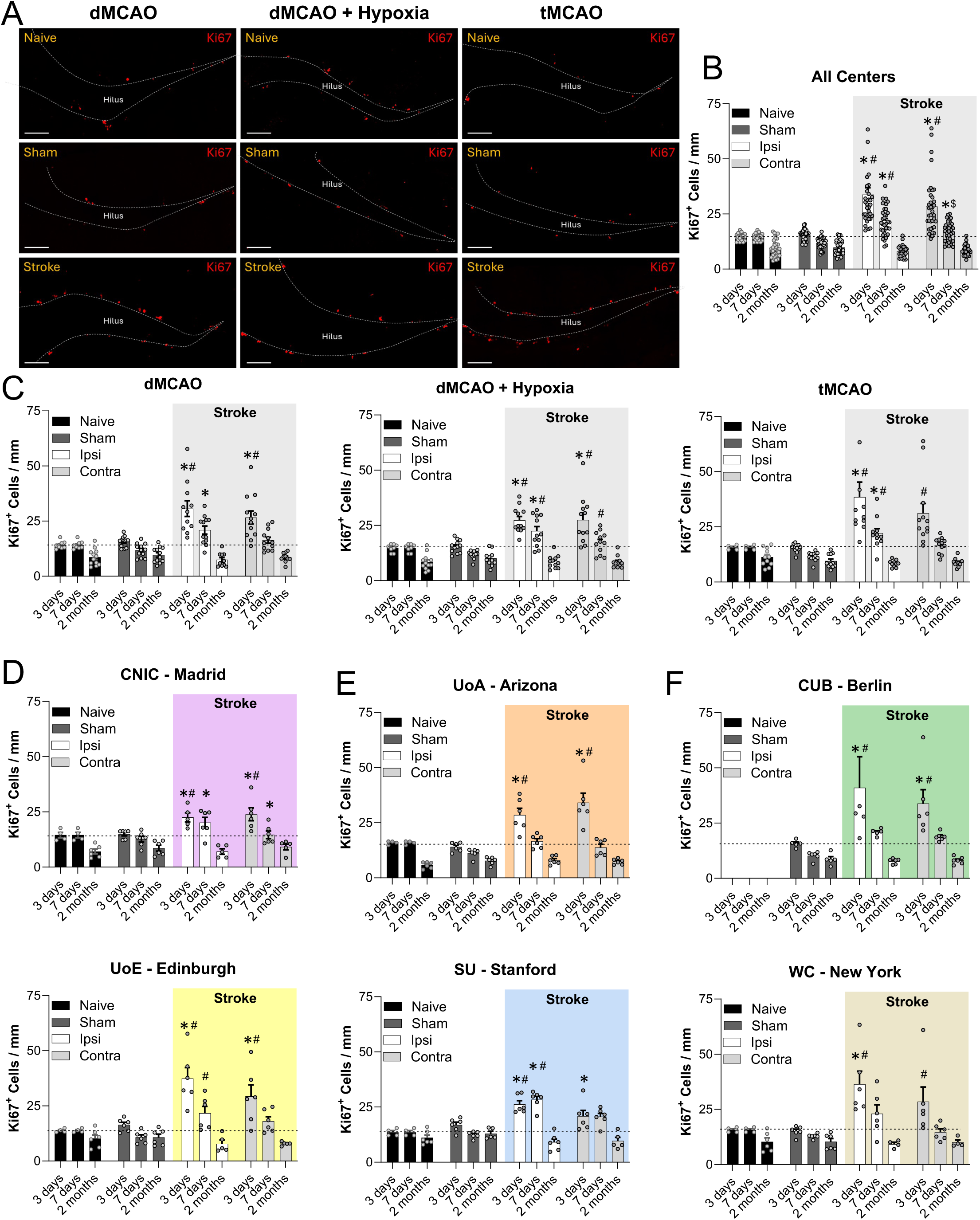
Multi-center analysis of post-stroke proliferation in the hippocampus. (A) Representative confocal images of Ki67⁺ proliferating cells (red) in the dentate gyrus from naive, sham and MCAO (ipsilateral) mice at 3 days post-stroke, illustrating examples from the three ischemia paradigms used (dMCAO, dMCAO + Hypoxia, and tMCAO). White dotted lines delineate the inner edge of the SGZ. Scale bar: 100 µm. (B) Quantification of Ki67⁺ cells per mm across all centers pooled, showing a robust bilateral increase in proliferation at 3 days and 7 days after stroke, returning to baseline by 2 months. (C) Ki67⁺ cell density grouped by ischemia model: permanent distal MCA occlusion (dMCAO; CNIC–Madrid and UoE–Edinburgh), permanent distal MCA occlusion combined with systemic hypoxia (dMCAO + hypoxia; UoA–Arizona and SU–Stanford), and transient proximal filament occlusion (tMCAO; CUB–Berlin and WC–New York). (D–F) Ki67⁺ cell quantification from individual centers with the corresponding ischemia paradigm indicated. (D) dMCAO centers: CNIC–Madrid (purple) and UoE–Edinburgh (yellow). (E) dMCAO + hypoxia centers: UoA–Arizona (orange) and SU–Stanford (blue). (F) tMCAO centers: WC–New York (beige) and CUB–Berlin (green). Bars represent mean ± SEM from 5–7 mice per group. Time points analyzed include 3 days, 7 days, and 2 months post-stroke (exact sampling varies by site). Groups shown are naive (black), sham (dark grey), ipsilateral stroke (white), and contralateral stroke (light grey). Statistical comparisons were performed using two-way ANOVA followed by Tukey’s post hoc test. *p<0.05 vs naive; ^#^p<0.05 vs sham; ^$^p<0.05 vs stroke ipsilateral.

### STROKE INDUCES A BILATERAL INCREASE IN NEUROBLASTS

We next assessed the density of DCX-positive neuroblasts at 7 days and 2 months after stroke **(Figure 3)**. At 7 days, a bilateral increase in DCX-positive cells was observed when all stroke groups were combined, with a similar response in both the ipsilateral and contralateral hippocampus **(Figure 3A-B).** This effect was significant compared with naive mice, indicating that ischemia robustly stimulates production and/or survival of immature neurons during the subacute phase. By 2 months, however, DCX levels no longer differed from baseline, suggesting that the neurogenic response does not persist long-term. Nevertheless, consistent with our previous findings^11^, we observed a significant expansion of the granule cell layer **(Supplementary Figure 1),** indicating that the early surge of neurogenesis leaves a lasting structural footprint in the dentate gyrus even after neuroblast numbers have normalized. Interestingly, sham-operated mice showed a mild elevation in DCX-positive cells in the ipsilateral side, with values falling between those of naive and ischemic animals. This intermediate profile suggests that the surgical procedure itself may induce a modest neurogenic response, likely reflecting tissue stress associated with the intervention. These patterns were generally consistent across the different ischemic paradigms (dMCAO, dMCAO + hypoxia, and tMCAO) as well as across individual sites, which showed similar trends despite methodological differences **(Figure 3C–F).**

**Figure 3.**
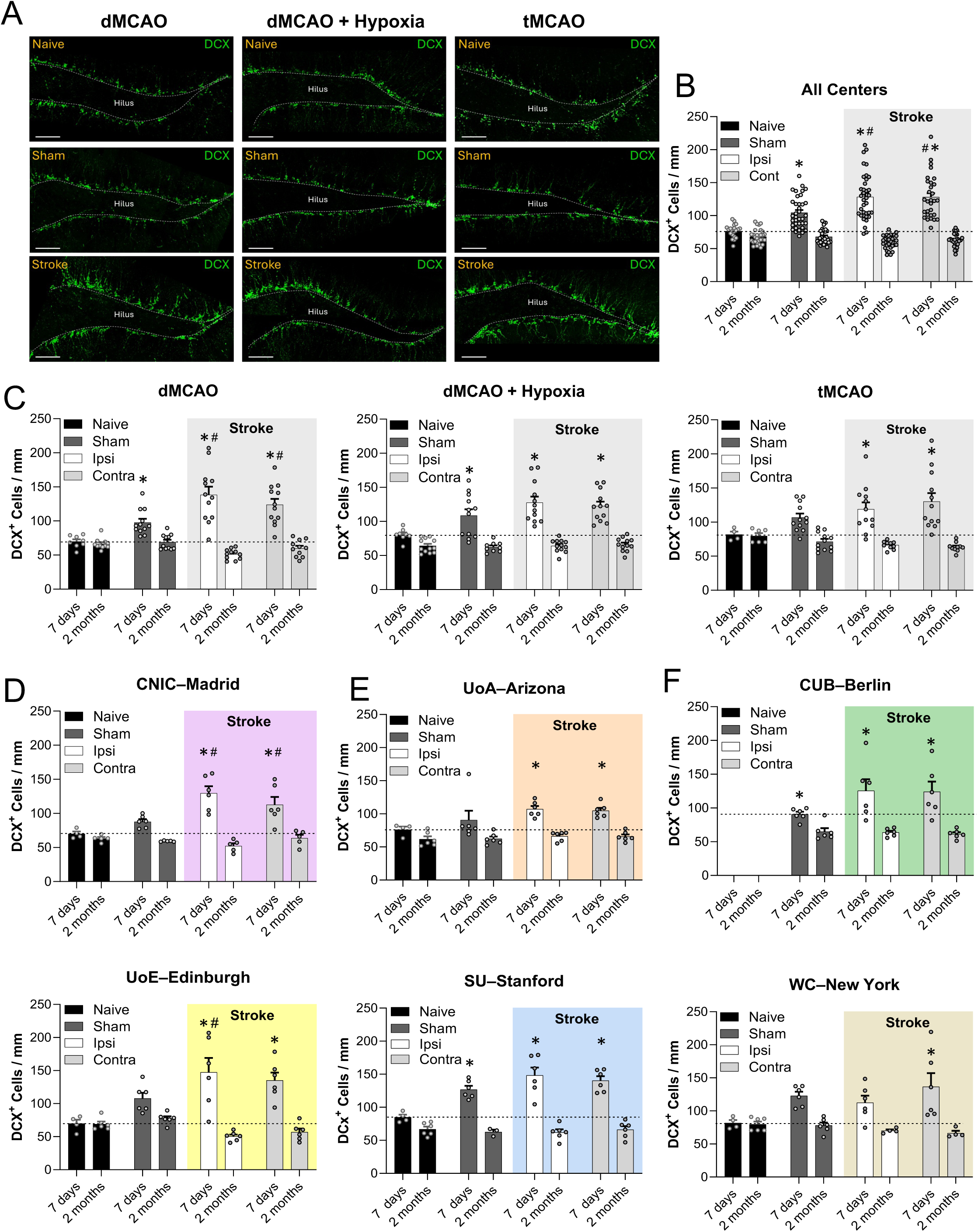
Multi-center analysis of post-stroke neuroblasts in the hippocampus. (A) Representative confocal images of DCX⁺ immature neurons (green) in the dentate gyrus from naive, sham and stroke (ipsilateral) mice at 7 days post-stroke, illustrating examples from the three ischemia paradigms used (dMCAO, dMCAO + Hypoxia, and tMCAO). White dotted lines delineate the inner edge of the SGZ. Scale bar: 80 µm. (B) Quantification of DCX⁺ cells per mm across all centers pooled, showing a robust bilateral increase in neuroblast density at 7 days post-stroke, normalizing by 2 months. (C) DCX⁺ cell density grouped by ischemia model: permanent distal MCA occlusion (dMCAO; CNIC–Madrid and UoE–Edinburgh), permanent distal MCA occlusion combined with systemic hypoxia (dMCAO + hypoxia; UoA–Arizona and SU–Stanford), and transient proximal filament occlusion (tMCAO; CUB–Berlin and WC–New York). (D–F) DCX⁺ cell quantification from individual centers with the corresponding ischemia paradigm indicated. (D) dMCAO centers: CNIC–Madrid (purple) and UoE–Edinburgh (yellow). (E) dMCAO + hypoxia centers: UoA–Arizona (orange) and SU–Stanford (blue). (F) tMCAO centers: CUB–Berlin (green) and WC–New York (beige). Bars represent mean ± SEM from ∼5–7 mice per group. Time points analyzed include 7 days and 2 months post-stroke. Groups shown are naive (black), sham (dark grey), ipsilateral stroke (white), and contralateral stroke (light grey). Statistical comparisons were performed using two-way ANOVA followed by Tukey’s post hoc test. *p<0.05 vs naive; ^#^p<0.05 vs sham.

### STROKE INDUCES ABERRANT FEATURES OF ADULT-GENERATED HIPPOCAMPAL NEURONS

Our previous work using a model of permanent ischemia induced by distal MCA ligation combined with common carotid artery occlusion^10,11^ demonstrated that post-stroke hippocampal neurogenesis contributes to cognitive impairment not only because more neurons are generated but also because a substantial proportion of them exhibit aberrant morphology. Therefore, we next analyzed the structural maturation of newborn neurons at 2 months post-ischemia. Based on our previous studies, we quantified apical dendrite (AD) length as a parameter of aberrant neurogenesis using DCX immunostaining at the 2-month time point. The pooled analysis across all centers revealed a significant shortening of the apical dendrite in newborn neurons in both the ipsilateral and contralateral hippocampus, when compared with both sham-operated and naive mice (**Figure 4A-B**). This effect became even more evident when neurons were classified into length categories (<15 µm, 15–40 µm, and >40 µm), revealing a marked shift toward the shorter-length classes in ischemic mice **(Figure 4C)**, with the distribution for each length interval presented separately in **Figures 4D**. Although sham-operated animals showed a mild increase in DCX-positive cells (**Figure 3**), only slight reductions in AD length were observed in shams at some centers (**Supplementary Figure 2**), consistent with modest effects of surgical manipulation on early neurogenic activity but insufficient to induce morphological abnormalities. Stroke-induced changes in apical dendrite length were dependent on both the ischemia paradigm and the post-stroke time point analyzed. At 2 months post-stroke, the most robust and consistent reduction in apical dendrite length was observed in the transient filament occlusion model (tMCAO), where significant bilateral shortening was detected across centers (**Figure 4E–F and Supplementary Figure 2**). In contrast, distal permanent occlusion models exhibited more variable outcomes at this chronic stage: while the UoE–Edinburgh cohort showed a significant reduction in AD length, no differences were detected in the CNIC–Madrid or UoA–Arizona cohorts, consistent with previous reports indicating dendritic shortening at earlier time points that is no longer evident at 2 months^11^. Together, these results suggest a time-dependent evolution of apical dendritic remodeling after stroke, with transient proximal occlusion leading to more persistent morphological alterations.

**Figure 4.**
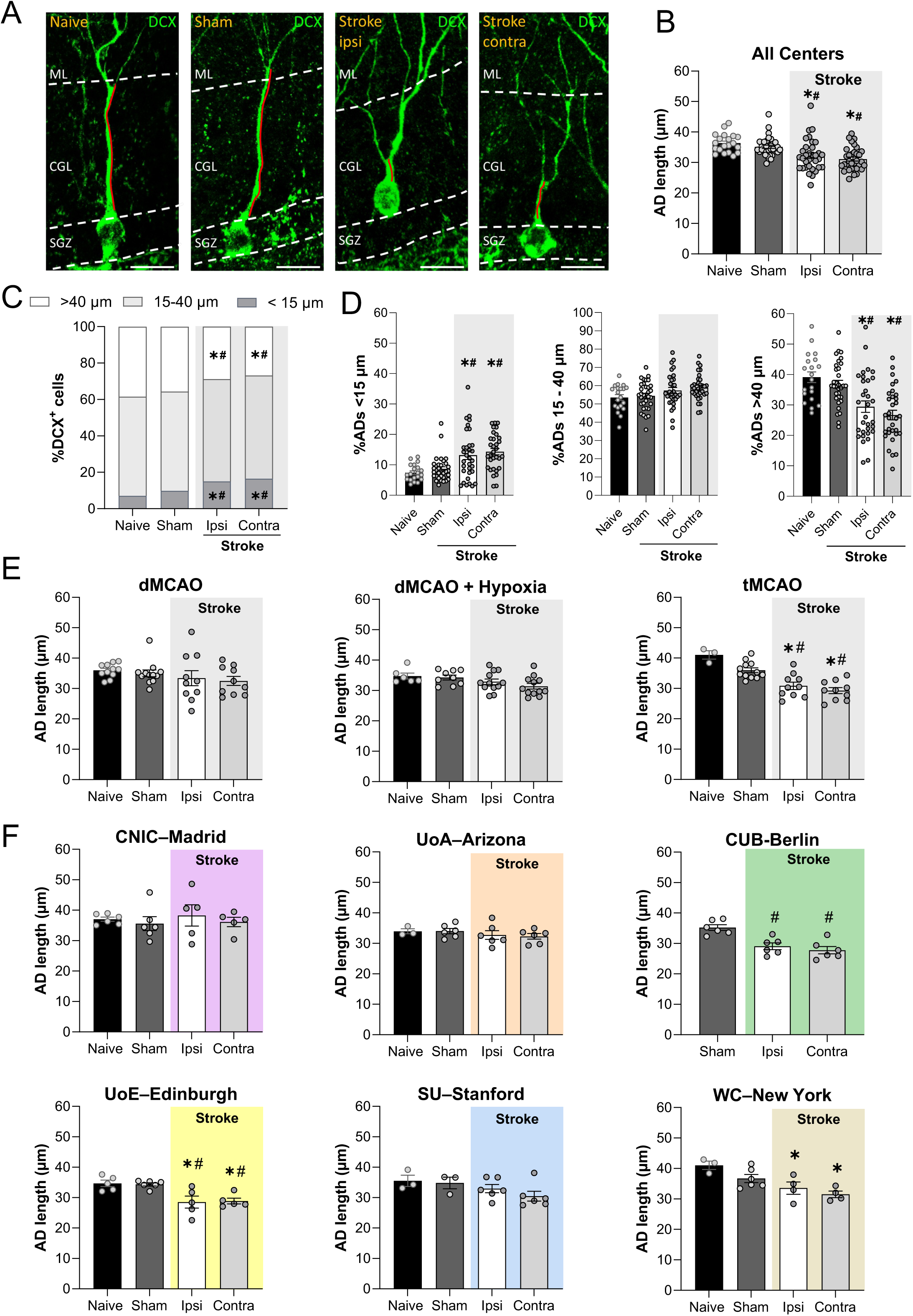
Analysis of neuroblast apical dendrite morphology in the dentate gyrus 2 months after stroke. (A) Representative confocal images from CUB–Berlin group of DCX⁺ immature neurons (green) from naive, sham and stroke mice (ipsilateral and contralateral hemispheres) illustrating apical dendrite (AD, red line) morphology. White dashed lines indicate the interface between molecular layer (ML), granule cell layer (GCL), and subgranular zone (SGZ). Scale bar: 10µm. (B) Quantification of apical dendrite (AD) length (µm) across groups shows a significant reduction in dendritic extension in both ipsilateral and contralateral hemispheres after stroke compared with naive and sham mice. (C) Stacked distribution of DCX⁺ neurons classified by AD length (<15 µm, 15–40 µm, >40 µm) demonstrating a shift toward shorter dendrites in stroke mice. (D) Bar graphs showing the percentage of DCX⁺ neurons within each AD length category (<15 µm, 15–40 µm, >40 µm) for naive, sham, ipsilateral stroke, and contralateral stroke groups. (E) AD length grouped by ischemia model: permanent distal MCA occlusion (dMCAO; CNIC–Madrid and UoE–Edinburgh), permanent distal MCA occlusion combined with systemic hypoxia (dMCAO + hypoxia; UoA–Arizona and SU–Stanford), and transient proximal filament occlusion (tMCAO; CUB–Berlin and WC–New York). (F) Quantification of AD length for each center individually. CNIC–Madrid (purple), UoE–Edinburgh (yellow), UoA–Arizona (orange), SU–Stanford (blue), CUB–Berlin (green), and WC–New York (beige). Bars represent mean ± SEM from approximately 5–7 mice per group, all analyzed at 2 months post-stroke. Data were analyzed using one-way ANOVA followed by Tukey’s post hoc test, or Kruskal–Wallis followed by Dunn’s post hoc test when appropriate. *p<0.05 vs naive; ^#^p<0.05 vs sham.

Finally, we quantified a range of aberrant neuronal morphologies previously reported in conditions where adult neurogenesis is disrupted, including inflammation, neurodegeneration, and stroke^9,11,21–23^, to determine how ischemia alters the maturation trajectory of newborn granule cells. These included ectopic neurons, multipolar neurons, laterally projecting neurons, and inverted neurons, each representing a distinct deviation from the canonical maturation of granule cells (**Figure 5A**). We incorporated a fifth category termed “aberrant” which encompassed all these morphological abnormalities. All five phenotypes were systematically evaluated in DCX-positive neurons at the 2-month time point (**Figure 5** and **Supplementary Figures 3-4**). In the pooled analysis across all centers, we observed a significant increase in the proportion of aberrant DCX⁺ neurons in ischemic mice compared with sham controls (**Figure 5B-C**). The percentage of aberrant cells nearly doubled after stroke, rising from approximately 2.2% in sham-operated animals to approximately 4.8-4.9% in both the ipsilateral and contralateral dentate gyrus. When the different morphological subclasses were examined individually, significant increases were detected in the proportion of multipolar (bipolar) cells and laterally grown neurons, whereas inverted and ectopic phenotypes did not differ significantly between groups (**Supplementary Figure 3A**). In this analysis, only sham-operated animals were used as controls; however, this does not affect interpretation because sham and naive animals do not differ in dendritic morphology as demonstrated in Figure 4, confirming that sham mice provide a valid baseline for assessing post-ischemic aberrant neurogenesis. When each ischemic paradigm was analyzed separately, all three models (dMCAO, dMCAO + hypoxia, and tMCAO) showed an increased proportion of aberrant DCX⁺ neurons compared with their respective shams **(Figure 5D)**. Across models, this increase was mainly driven by the multipolar/bipolar and laterally grown categories, whereas inverted and ectopic phenotypes remained unchanged **(Supplementary Figure 3B–D)**, indicating that stroke consistently affects specific aspects of newborn neuron maturation. Finally, site-specific analyses further confirmed these patterns, with individual centers showing comparable distributions of aberrant morphologies despite methodological differences, underscoring the robustness and reproducibility of these findings across laboratories (**Figure 5E and Supplementary Figure 4**).

**Figure 5.**
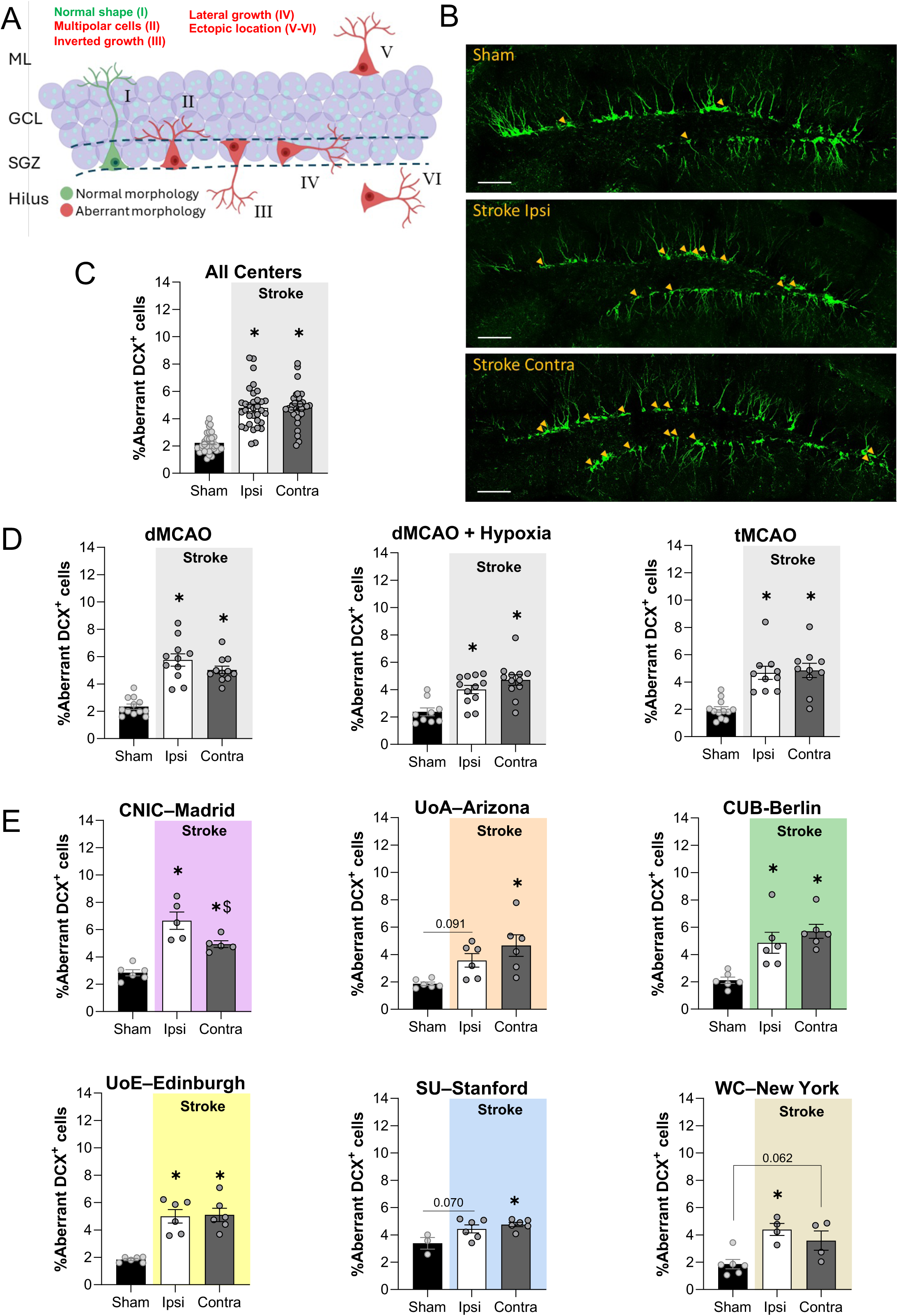
Analysis of aberrant DCX⁺ neuroblast morphology in the dentate gyrus at 2 months post-stroke. (A) Schematic representation of normal and aberrant DCX⁺ immature neurons in the dentate gyrus, illustrating the five morphological categories analyzed: normal Y-shaped morphology (I), multipolar cells (II), inverted somatodendritic polarity (III), lateral growth (IV), and ectopic localization within the upper granule cell layer or hilus (V–VI). (B) Representative confocal images from CUB–Berlin group of DCX⁺ neuroblasts (green) from sham, ipsilateral stroke, and contralateral stroke hemispheres highlighting aberrant morphologies (yellow). Scale bars: 100 μm. (C) Pooled quantification across all centers showing the percentage of DCX⁺ neurons displaying any aberrant morphology in sham, ipsilateral, and contralateral stroke hemispheres. (D) Model-specific quantification of total aberrant DCX⁺ neurons for permanent distal MCA occlusion (dMCAO), dMCAO combined with hypoxia, and transient filament MCA occlusion (tMCAO). (E) Center-level analyses showing the percentage of aberrant DCX⁺ neurons for each participating site and corresponding ischemia model: CNIC–Madrid and UoE–Edinburgh; SU–Stanford (dMCAO) and UoA–Arizona (dMCAO + hypoxia); and WC–New York and CUB–Berlin (tMCAO). Bars represent mean ± SEM from 5–7 mice per group. Statistical comparisons were performed using one-way ANOVA with Tukey’s post hoc test or Kruskal–Wallis with Dunn’s post hoc test when appropriate. *p<0.05 vs sham; ^$^p<0.05 vs stroke ipsilateral. Individual aberrant morphology categories (multipolar, lateral, inverted, ectopic) are presented separately in Supplementary Figure 3 and 4. Schematic illustration in panel A was created with BioRender.

## DISCUSSION

Stroke triggers a profound remodeling of AHN, yet whether this response is conserved across experimental paradigms has remained uncertain. By integrating data from six independent centers using multiple ischemia models, we show that the hippocampal neurogenic response follows a remarkably stable pattern despite substantial differences in vascular territory, reperfusion status, surgical approach, inflammatory context, and laboratory environment. This consistency indicates that key features of post-stroke neurogenesis are largely model-independent and reflect fundamental properties of the post-ischemic hippocampal niche.

Across all models, we observed a robust bilateral increase in SGZ proliferation at 3 days that persisted, albeit at reduced magnitude, at 7 days. DCX⁺ neuroblast density followed a similar trajectory, increasing consistently across paradigms without association with injury severity or spatial distribution. Importantly, although tMCAO produced the largest infarcts and caused hippocampal cell loss in a subset of animals^13^, its proliferative and neuroblast responses were comparable to those of permanent distal occlusions with or without hypoxia, which typically generate smaller and cortex-restricted lesions. These findings indicate that hippocampal activation does not scale with infarct size nor require direct hippocampal injury. Instead, progenitor recruitment is likely driven by global post-ischemic disturbances such as hypoperfusion, metabolic and oxidative stress, neurotransmitter imbalance, immune dysregulation, and network instability, that are shared across ischemic insults^24,25^. The uniformity of this early response raises important questions about its long-term consequences. A rapid surge in hippocampal progenitor activation followed by apparent normalization may reflect resolution of acute stress, but it may also impose lasting strain on the neurogenic niche. Evidence from aging, chronic inflammation, traumatic brain injury and seizures suggests that repeated or intense activation of radial glia–like (RGLs) stem cells can compromise their self-renewal capacity or bias their lineage output^26–28^. Although stem cell populations were not directly assessed here, the strikingly similar proliferative response across models suggests that ischemia may exert comparable pressure on the hippocampal stem cell compartment. Future studies should determine whether stroke alters the quiescence-activation balance of RGLs, affects their long-term proliferative potential, or drives lineage bias toward aberrant neuronal maturation.

A central finding of this multi-center analysis is that normalization of proliferation and DCX⁺ cell numbers by 2 months does not signify restoration of the neurogenic process. Despite quantitative recovery, newborn neurons exhibited persistent and characteristic morphological abnormalities, including alterations in somatodendritic polarity, ectopic or lateral growth, and, at this chronic time point, truncation of the apical dendrite in a model-dependent manner. While some of these features were observed across ischemia paradigms and centers, apical dendrite shortening at 2 months post-stroke was most prominent and consistent following transient proximal occlusion. Together, these findings indicate that the neurogenic lineage undergoes durable reprogramming toward an aberrant maturation trajectory that extends well beyond the acute phase, even in the absence of sustained changes in cell number.

These results extend previous single-laboratory studies demonstrating that aberrant neurogenesis contributes causally to long-term hippocampus-dependent cognitive impairment^10,11^ and that suppressing post-stroke neurogenesis prevents chronic cognitive deficits. Our prior work showing that increased inhibitory tone in the ipsilesional hippocampus correlates with both aberrant neurogenesis and cognitive decline further supports a mechanistic role for dysfunctional neuronal maturation rather than insufficient neurogenesis^10^. The present multi-center dataset provides strong convergent evidence that neurons generated after stroke are structurally unfit for proper circuit integration and that impaired maturation persists months after injury.

Building on the observation that early proliferative and neuroblast responses were remarkably uniform across ischemia paradigms and independent of infarct size or distribution, we next examined whether long-term morphological outcomes followed a similar pattern. Despite marked differences in lesion extent and, in some cases, direct hippocampal involvement in the tMCAO model^13^, the spectrum of aberrant neuronal phenotypes observed at later stages did not segregate according to injury severity or spatial proximity. Instead, alterations in neuronal polarity, positioning, and structural maturation were detected across ischemia paradigms, indicating that maladaptive neurogenesis reflects a common downstream consequence of ischemia rather than a direct effect of focal tissue loss. With respect to AD length, our data indicate a clear temporal component. In previous work, significant apical dendrite shortening was detected one month after ischemia in the Madrid dMCAO model, whereas at 2 months post-stroke this phenotype was no longer evident, suggesting partial structural recovery over time. In contrast, in the present multi-center analysis, significant bilateral AD shortening persisted at 2 months in the tMCAO cohorts and in the UoE–Edinburgh dMCAO cohort, indicating that the durability of dendritic alterations differs across ischemia paradigms and sites. Importantly, however, AD length alone did not capture the full extent of neurogenic disruption. Rather, the sustained presence of ectopic, misoriented, and structurally immature granule neurons, reflected in persistent polarity defects and elevated proportions of aberrant cells, emerged as the most robust and functionally relevant hallmark of post-stroke hippocampal remodeling. Thus, even when apical dendrite length partially normalizes at later stages, maladaptive neurogenesis remains a durable consequence of cerebral ischemia.

Sham-operated animals further reinforced the specificity of the ischemic response. Although shams occasionally showed mild, transient increases in proliferation or DCX expression at early time points, likely reflecting surgical manipulation, anesthesia, perioperative inflammation, or, in paradigms involving systemic hypoxia, exposure to hypoxic conditions in the absence of focal ischemia, these effects did not translate into overt long-term structural alterations. At 2 months post-surgery, sham and naive animals exhibited broadly comparable hippocampal cytoarchitecture, with no granule cell layer enlargement and only modest changes in dendritic morphology. Notably, sham animals exposed to hypoxia displayed a higher proportion of aberrant dendritic phenotypes than other sham groups, yet these changes remained substantially less pronounced than those observed after stroke. Crucially, shams did not develop the full spectrum of ectopic positioning, polarity defects, and persistent structural immaturity characteristic of post-ischemic neurogenesis, confirming that the abnormalities identified here primarily reflect consequences of the ischemic insult rather than surgical artefacts.

Several methodological strengths support the robustness of these conclusions. This is, to our knowledge, the first multi-center, multimodel study examining hippocampal neurogenesis after stroke. By integrating data from six independent laboratories employing distinct MCAO approaches - including transient filament occlusion, permanent distal occlusion by ligation or cautery, and permanent occlusion combined with hypoxia- we capture a wide spectrum of ischemic pathophysiology. Harmonized analysis pipelines, blinded quantification, and large cumulative sample size further enhance reproducibility and translational relevance. Moreover, the inclusion of both early (proliferation and neuroblast density) and late (morphology and positional integration) neurogenic endpoints provides a comprehensive view of post-stroke neurogenic trajectories.

Several limitations should be acknowledged. The exclusive use of male mice precludes assessment of sex-specific responses. Behavioral and electrophysiological analyses were not incorporated, preventing direct correlation between structural abnormalities and functional outcomes. Although multi-center designs inevitably introduce variability, the strong qualitative convergence across sites mitigates concerns about site-specific artefacts. Additionally, all experiments were conducted in young adult mice, whereas stroke predominantly affects older individuals. Because hippocampal neurogenesis declines markedly with age^29–31^ and aging reshapes progenitor quiescence, inflammatory tone, and neuronal maturation^26,32^, neurogenic responses to ischemia may differ in older brains. Future studies incorporating aged cohorts will be essential. Finally, we did not directly evaluate RGL stem cells or upstream molecular mechanisms, leaving open questions regarding long-term stem cell depletion, lineage bias, and the pathways sustaining maladaptive neurogenesis. Moreover, other neurogenic regions, such as the SVZ, may also contribute to chronic post-stroke outcomes and warrant analogous multi-center investigation.

In summary, these findings have important translational implications. They challenge the long-standing assumption that stimulating neurogenesis is inherently beneficial after stroke and instead identify maladaptive neuronal maturation as a key contributor to chronic cognitive impairment. Given that at least 30–35% of stroke survivors develop persistent cognitive deficits or dementia^33–36^, strategies aimed at correcting or preventing abnormal maturation of newborn neurons may represent a more effective therapeutic avenue. Potential targets may include inhibitory circuit balance, microglial activation, neurotrophic signaling, or extracellular matrix composition. The conserved morphological abnormalities identified here also provide robust preclinical readouts for therapeutic screening. By demonstrating that aberrant hippocampal neurogenesis is a model-independent hallmark of cerebral ischemia, this study lays the groundwork for future interventions designed to preserve the quality, rather than the quantity, of adult-born neurons after stroke.

## DATA AND MATERIALS AVAILABILITY

The data are presented in the main manuscript and in the supplementary materials.

## ACKNOWLEDGMENTS

We thank all members of our laboratory and the Stroke-IMPaCT consortium for their insightful feedback and collaborative support; E. Garrido and S. Rodriguez for animal husbandry; E. Arza, O. Giménez, V. Labrador, V. Caiolfa from the Microscopy Unit of the CNIC for help with microscopy.

